# Metagenomic enables the first detection of *Trypanosoma* sp. in *Streblidae* (Diptera: *Hippoboscoidea*) parasitizing bats in São Paulo, Brazil

**DOI:** 10.1101/2025.06.12.659397

**Authors:** Roberta Marcatti, Lucas Augusto Moysés Franco, Esmênia Coelho Rocha, Marcello Schiavo Nardi, Juliana Laurito Summa, Eric Thal Brambilla Cordeiro da Silva, Adriana Ruckert da Rosa, Débora Cardoso de Oliveira, Gustavo Graciolli, Ester Cerdeira Sabino

## Abstract

Bats play important ecological roles but also can carry various pathogens, including trypanosomatids. This study reports the first detection of *Trypanosoma* sp. in flies from the Streblidae family parasitizing *Carollia perspicillata* bats in a peri-urban fragment of the Atlantic Forest in São Paulo. Using shotgun metagenomics, we detected Trypanosoma sequences in half of the fly samples analyzed. Phylogenetic analysis showed these sequences belong to the Neobat 4 clade, previously reported in *Carollia* sp. bats. Although streblid flies’ ability to transmit Trypanosoma is still unknown, their close connection with bats means they might be used as a non-invasive way to monitor pathogens when bat sampling is difficult. More research is needed, but this work expands what we know about the Neobat 4 clade’s geographic distribution and parasite circulation in bats and their ectoparasites.

## INTRODUCTION

Bats are flying mammals distributed on all continents except Antarctica and represent around 20% of all mammal species, with over 1,400 known species in total and their origin is estimated to have occurred during the Eocene, around 60 million years ago [1]. They provide numerous ecosystem services and their preservation is extremely important for the maintenance of green areas [2]. Despite their undeniable ecological importance, bats are also considered hosts of numerous infectious agents of importance to Public Health, such as the rabies virus, coronavirus and paramyxovirus, highlighting the need of continuous pathogen surveillance to better understand the spatial distribution of these agents in natural areas, especially within the current scenario of intense climate change [3].

These mammals have an intrinsic relationship with a hematophagous ectoparasite specific to this animal group: dipterans of the Families Streblidae and Nycteribiidae. Although both have evolved in close association with bats, specific co-evolution with host species has resulted in distinct morphological adaptations for each group. This close relationship and the resulting adaptations make these dipterans excellent models for the study of host-parasite co-evolution [4]. These flies show high host specificity and strategies that maximize reproduction, including host switching within colonies [5–7], and authors suggest these characteristics are key factors in pathogen transmission dynamics [8]. The Streblidae family includes about 230 species, which are mainly found in tropical regions [9].

Bats are natural reservoirs of trypanosomatids from the cruzi clade and possibly contributed to their dispersal between continents [10]. The Trypanosomatidae family includes the genera *Trypanosoma, Leishmania*, and *Crithidia*, which are known to cause diseases in various hosts [11]. The genus *Trypanosoma* causes two diseases of great medical relevance: Chagas disease (*Trypanosoma cruzi*) and African sleeping sickness (*Trypanosoma brucei*). Transmission occurs mainly through hematophagous invertebrate vectors, with triatomines, popularly known as kissing bugs, being the main vector [9].

In Brazil, infections by *T. cruzi, T. rangeli, T. c. marinkellei, T. dionisii*, and *T. wauwau* have been identified in bats [12,13]. To date, there have been no reports of trypanosomatid detection in streblid flies.

This strong dependence on bats, considered hosts of several infectious agents, makes these flies serve as sentinels. For this reason, an exploratory metagenomic study in bat flies was conducted, since this Next Generation Sequencing molecular technique allows the identification of several microorganisms contained in a sample, without the need for prior knowledge of the agents of interest.

The objective of this study is to report the first detection and preliminary genetic characterization of *Trypanosoma* sp. in Streblidae flies parasitizing bats, contributing to knowledge about the parasite’s distribution in peri-urban ecosystems in São Paulo municipality.

## MATERIALS AND METHODS

### Study Area

Samples were collected in August 2022 at the Anhanguera Wildlife Refuge (RVS), located in the Perus district, North Zone of São Paulo (Figure 1), approximately 38 km from the city center (lat. -23° 47’ 57.69” S, long. -46° 40’ 45.24” W). With about 800 hectares, the area is part of the Northern Atlantic Forest Ecological Corridor and is a priority in the Municipal Plan for Atlantic Forest Conservation and Recovery - PMMA São Paulo.

**Figure 1.**
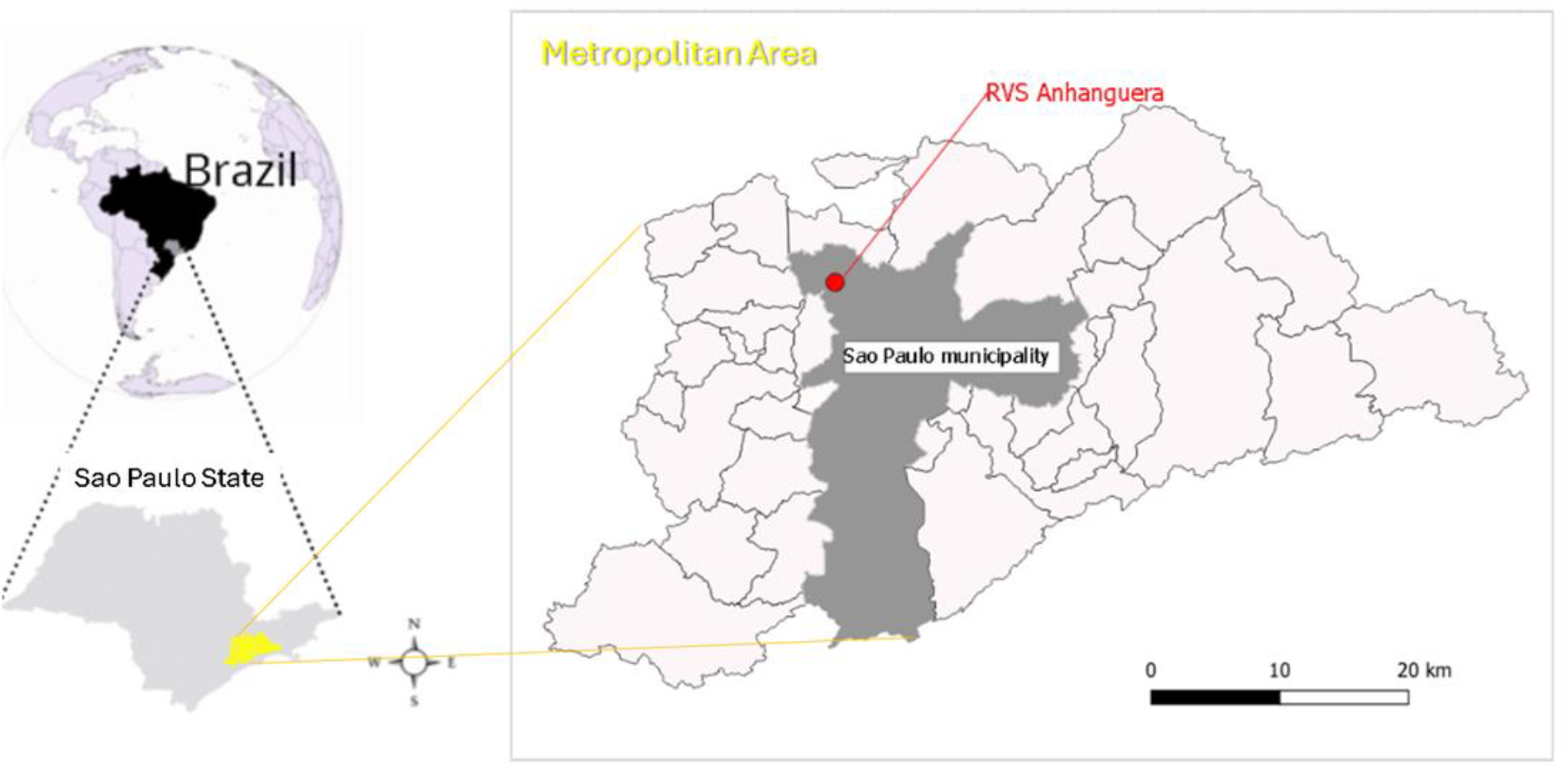
Location map of the metropolitan region and municipality of São Paulo, with study area (RVS Anhanguera) highlighted in red.

### Capture and Sample Preparation

Bats were captured using 10-meter-long mist nets with 50mm mesh, set up at dusk and checked every 30 minutes for 5 hours each night. *Streblidae* flies found parasitizing bats were collected and stored in cryotubes at -80°C, following the same identification as their host, processed on a cold table at -80°C for species identification. The specimens (bats and flies) were identified to species level. After pre-treatment to remove external contaminants with 70% ethanol for 10 seconds and three rinses in DNA/RNA-free water, the specimens were macerated with 350 µL of phosphate-buffered saline (PBS) and 3mm stainless steel beads in the L-beader 6 cell disruptor (Loccus, São Paulo, Brazil). After centrifugation at 10000 RPM for 10 minutes at 19ºC, the supernatant was processed through a 0.2-micron filter (Merck, Darmstadt, Germany). For every 150µL of sample, 7.5 µL of Proteinase K (Loccus, São Paulo, Brazil) was added, followed by incubation at 37º for 10 minutes in a dry bath. DNA and RNA extraction was performed using Extracta - DNA and RNA Viral (Loccus, São Paulo, Brazil), following the manufacturer’s protocol.

### Metagenomics Protocol

The SMART-9N protocol [14] was used for metagenomic sequencing. In brief, genetic material extracted from samples was treated with Turbo-DNase (Thermo Fisher Scientific, Massachusetts, USA), purified using the Zymo RNA Clean-up & Concentrator-5 kit (Zymo Research, California, USA), and cDNA was synthesized using SuperScript IV (Thermo Fisher Scientific, Massachusetts, USA). Amplification was performed with Q5 Hot Start High-Fidelity 2X Master Mix (New England Biolabs Massachusetts, USA). Sequencing libraries were prepared using the Native barcoding V14 kit (Oxford Nanopore Technologies, Oxford, UK) and sequenced on GridION (Oxford Nanopore Technologies, Oxford, UK) using the R10.4.1 flow cell (Oxford Nanopore Technologies, Oxford, UK). Raw data were converted to FASTQ for bioinformatic analysis.

### Bioinformatics

Raw data were processed in Genious Prime software (version 2024.1.1), with trimming via BBDuk (version 39.06) and removal of human and ectoparasite genome sequences. The remaining reads, longer than 500 bp, were taxonomically classified, with validation by BLASTn at NCBI (query cover > 70%) and the species of Trypanosoma that showed the highest similarity to the assembled reads was selected for the final alignment step.

Descriptive statistical analyses were conducted, and relative frequencies were calculated to assist in classifying the obtained sequences at the species level.

For contig assembly, Tadpole (BBTools) (version 39.06) was used, and alignment to the 18S rRNA region was performed with Minimap2 (version 2.26). This region, located in the small subunit of the ribosome, is frequently utilized for phylogenetic studies and species identification.

Phylogenetic inference was performed using MEGA12 software in two steps. First, a Maximum Likelihood tree (Tree 1) was built with 1000 bootstrap replications. The TN93+G evolutionary model (Tamura-Nei with Gamma distribution) was selected as the most suitable based on the Bayesian Information Criterion (BIC) applied to the data. This tree included sequences obtained in this study and previously described Trypanosoma species sequences from the T. cruzi clade (Figure 2).

**Figure 2.**
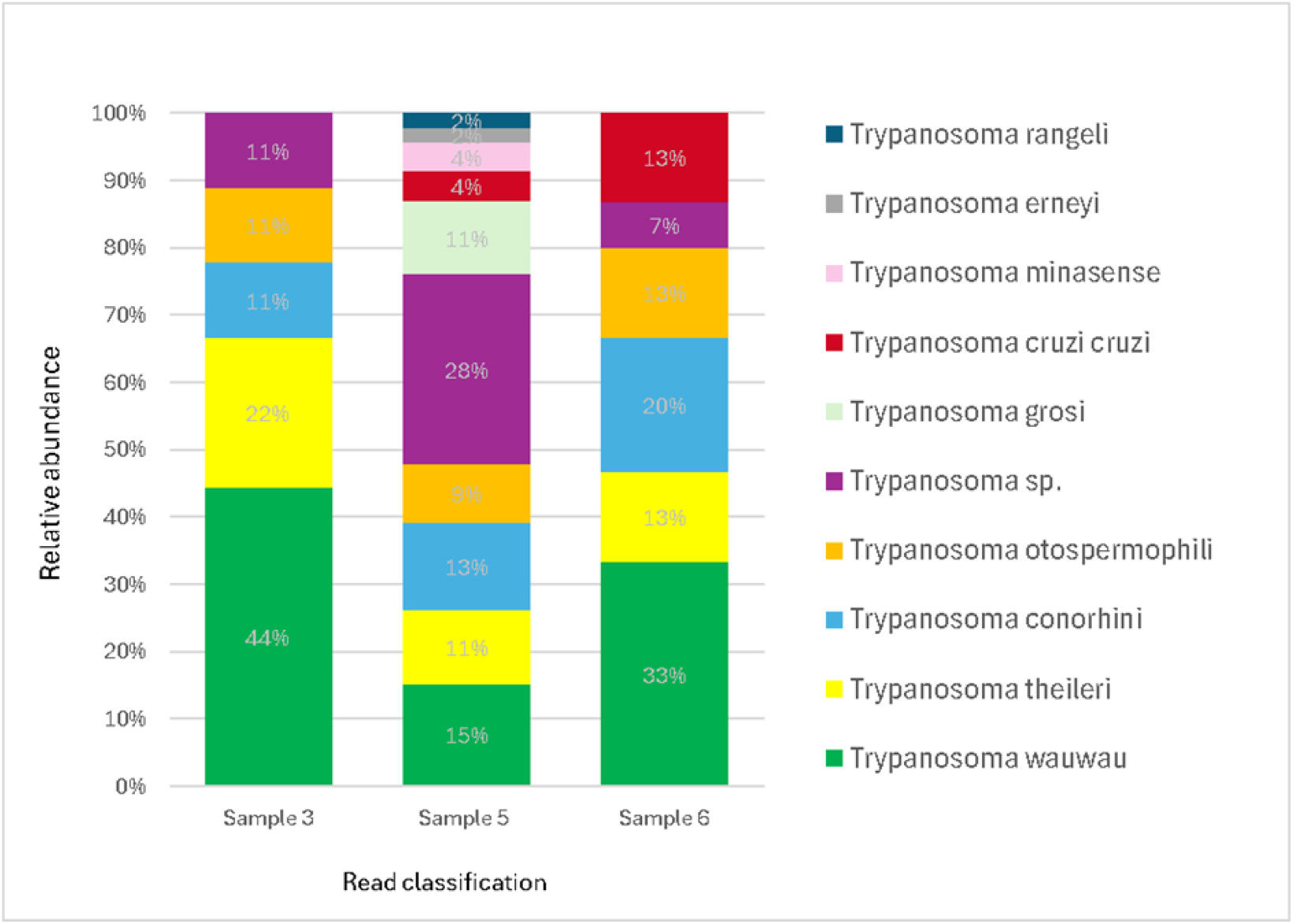
Relative abundance of read classification, per sample of Streblidae flies.

In the second stage, an additional phylogenetic tree (Tree 2) was constructed to analyze the distribution of generated *Trypanosoma* sp. sequences in relation to other unclassified species sequences obtained from bats in previous studies. This tree was generated using the Maximum Likelihood method, with 1000 bootstrap replicates and the TNe+I evolutionary model (Tamura-Nei with invariable sites proportion). This strategy was used to explore the diversity and evolutionary relationships of bat-associated *Trypanosoma* species in Brazil (Figure 3).

**Figure 3.**
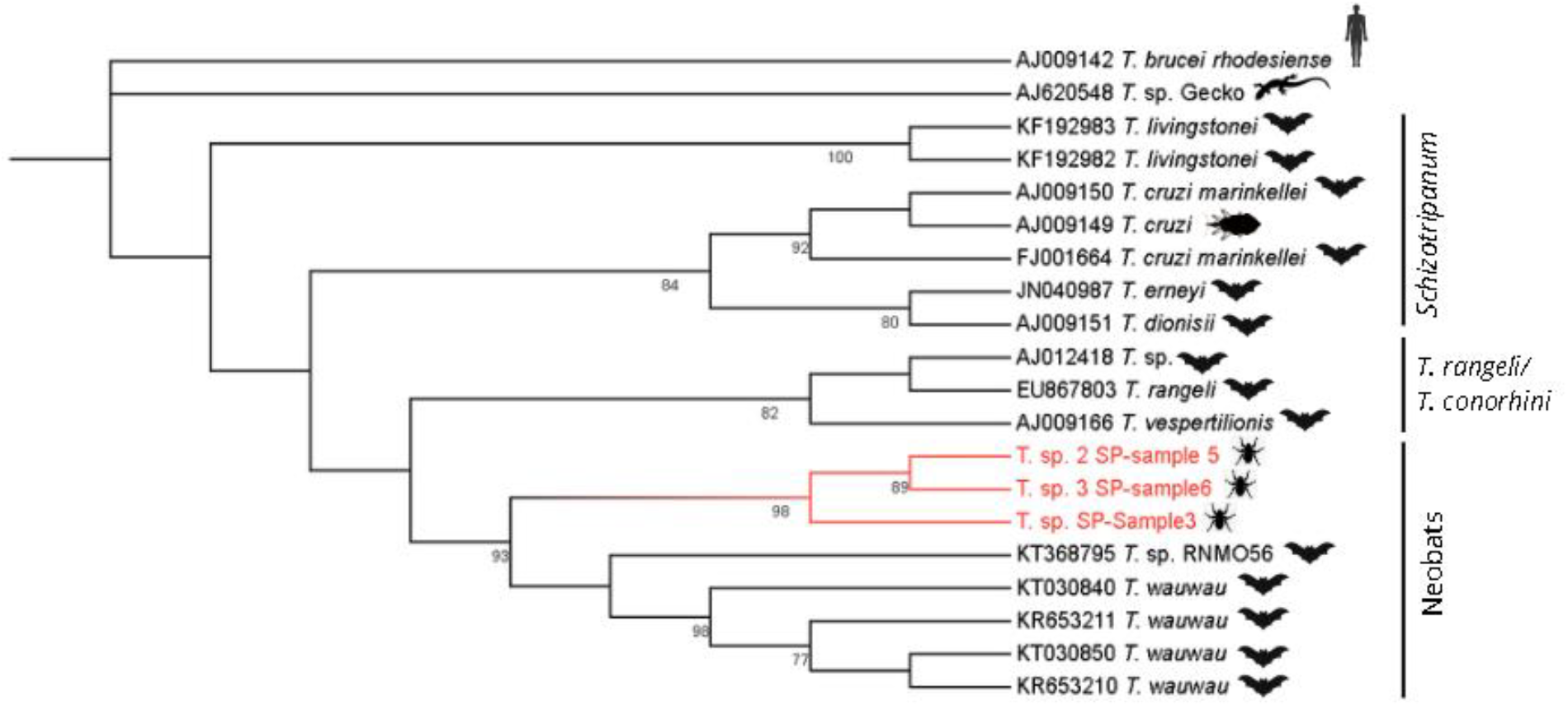
Phylogenetic tree of *Trypanosoma* spp. based on 18S SSU rRNA subunit sequences obtained by metagenomics of *Strebla guajiro* (Streblidae), with samples 3, 5 and 6 highlighted in red located in the Neobats clade.

The samples analyzed were collected during a single day, selected based on prior metagenomic findings that indicated the presence of trypanosomes. Samples from other days, in which no trypanosome sequences were detected, were not included in this analysis. As this is a preliminary result with a small sample size, no statistical association tests were performed with any other variables of interest.

### Ethics Approval and Consent to Participate

The project was approved by the Ethics Committee of the USP Faculty of Medicine (protocol No. 1439/2020), with a license for animal capture issued by the Secretariat of the Environment and Green Areas, São Paulo City Hall, SP, Brazil, responsible for wildlife management in the municipality.

## RESULTS

Among the 19 bats captured on the sampling day, six were parasitized with Streblidae (31.6%). Analysis of the six specimens revealed that three of them contained sequences of the genus *Trypanosoma* (Table I).

**Table 1.** Bat species captured, their respective ectoparasitic flies (Streblidae) and number of *Trypanosoma* reads detected by metagenomics.

| Bat species | Sex | Age | Streblidae species<br>(n especimens) | n *Trypanosoma*<br>sp. Reads |
| --- | --- | --- | --- | --- |
| *Anoura caudifer* | M | A | *Trichobius tiptoni* (1) | - |
| *Carollia perspicillata* | F | A | *Paraeuctenodes similis* (1)<br>*Trichobius joblingi* (2) | - |
| *Carollia perspicillata* | F | A | *Strebla guajiro* (1) | 388 |
| *Carollia perspicillata* | F | A | *Trichobius joblingi* (1) | - |
| *Carollia perspicillata* | F | A | *Strebla guajiro* (1) | 2448 |
| *Carollia perspicillata* | F | A | *Strebla guajiro* (1) | 317 |

The reads used for taxonomic classification are described in Table II. Among the three samples positive for *Trypanosoma*, samples “3” and “6” showed 20 reads larger than 500 bp, while sample “5” had approximately seven times more reads above 500 bp. For read homogeneity, 60 reads from sample “5” and 20 reads from each of the other samples were selected for BLASTn (Megablast) analysis. Only reads with query cover above 70% were considered for species-level classification.

**Table 2.** Absolute number of *Trypanosoma* reads detected by metagenomics.

| Sample | Reads<br>*Trypanosoma* | Reads above<br>500 bp | Reads above<br>70% coverage | Read Average<br>size (bp)* |
| --- | --- | --- | --- | --- |
| 3 | 387 | 20 | 9 | 618 |
| 5 | 2448 | 189 >>>> 60 | 46 | 620 |
| 6 | 319 | 20 | 15 | 611 |

\*Size of reads that were greater than 70% coverage

The normalized results by relative frequency (Fig. 2) showed the predominance of *Trypanosoma wauwau* in all three samples, with frequencies of 44%, 15%, and 33% respectively. This species was used as a reference for the final sequence assembly, focusing on the 18S rRNA region of the ribosomal gene.

Using the sequence assembly approach, we recovered fragments of 534 bp (*sample3*), 823 bp (*sample5*), and 814 bp (*sample6*). In the phylogenetic analysis, the sequences from this study formed a monophyletic clade with high bootstrap support (98%), positioning evolutionary closer to *Trypanosoma wauwau* than to other species belonging to the *Schizotrypanum* group clade (Figure 3). Bootstrap values above 70 are indicated at the nodes, while sequences from this study are highlighted in red. *T. brucei* and lizard *Trypanosoma* sequences were used as outgroups.

The second phylogenetic tree (Fig. 4), containing only bat *Trypanosoma* spp., incorporated the findings of Alves et al. [15] regarding uncultivable *Trypanosoma* sp. sequences, which described different Molecular Operational Taxonomic Units (MOTUs), classifying them as Neobat 1, 2, 3, 4, and 5. Other Brazilian *Trypanosoma* sp. sequences available in the literature were also included to determine the alignment of the new sequences with the clades proposed by Alves et al. *Trypanosoma brucei rhodesiense* and *T*. sp. from gecko lizard were used as outgroups.

**Figure 4.**
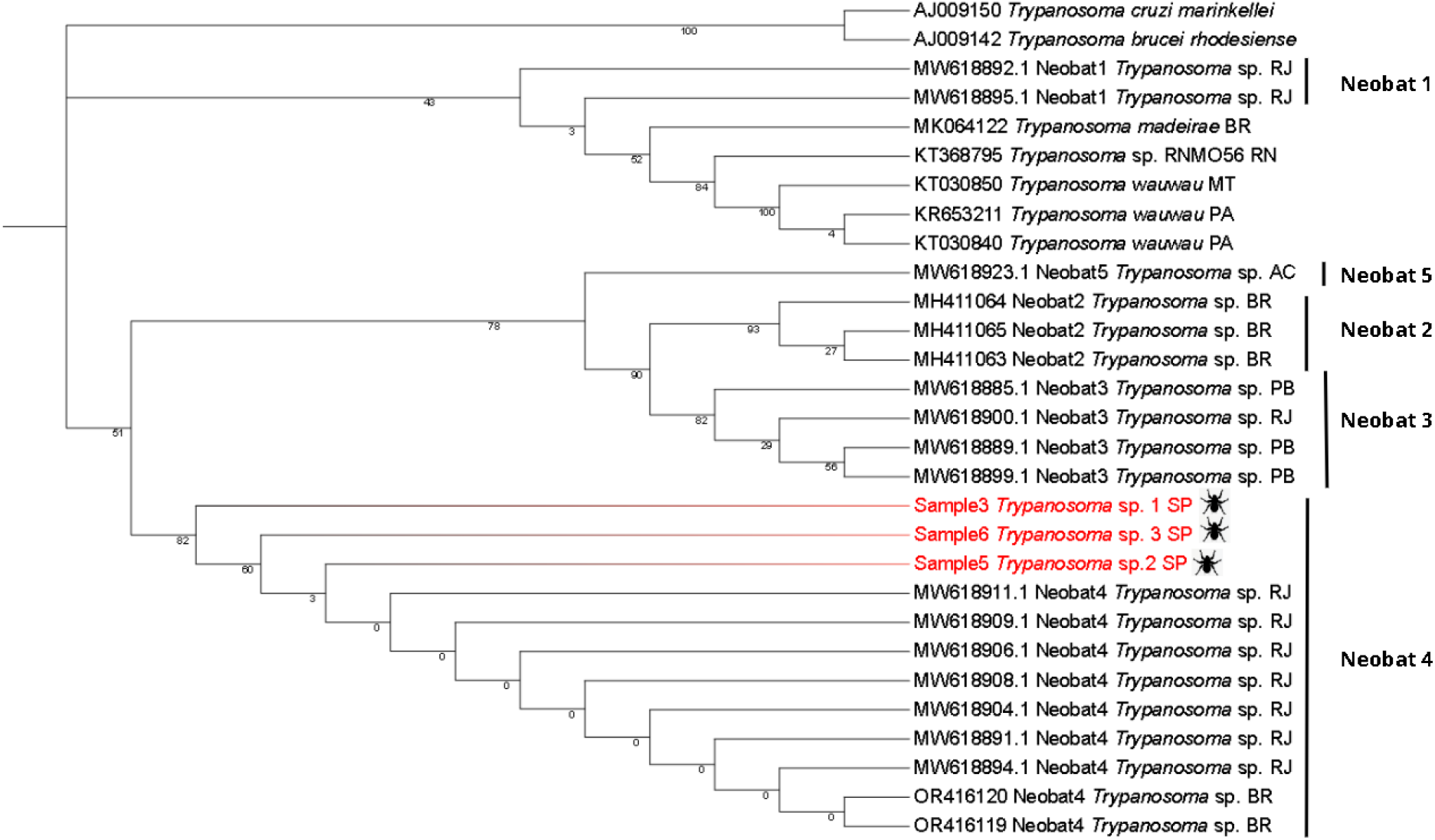
Phylogenetic tree containing *Trypanosoma* sp. sequences, based on the 18S SSU rRNA region obtained by metagenomics of *Strebla guajiro* (Streblidae), with samples 3, 5 and 6 highlighted in red. Sequences are indicated with Accession number, clade classification and geographic location in Brazil: Rio Grande do Norte (RN), Brazil without specified location (BR), Pará (PA), Mato Grosso (MT), Rio de Janeiro (RJ), Acre (AC), Paraíba (PB) and São Paulo (SP).

Our phylogenetic analysis confirmed the robustness of the main Neobat groupings, with significant bootstrap values of 0.9 for the Neobat2 clade and 0.8 for the relationship between Neobat2 and Neobat3, supporting the structure of 5 distinct lineages. The Neobat4 clade shows an interesting pattern in bootstrap values, with robust statistical support at basal nodes (0.823 and 0.601), but considerably low values in internal branches (0.003). Regarding geographic distribution, the Neobat4 clade is present in Rio de Janeiro and, with the three new sequences identified, in São Paulo. Neobat3 shows a broader distribution, including samples from both Paraíba and Rio de Janeiro. Available data in GenBank (NCBI) did not reveal geographic location information for sequences belonging to the Neobat2 clade, indicating a possible gap in deposited metadata.

## DISCUSSION

The detection of *Trypanosoma* in this study was possible by the untargeted nature of metagenomics, since this parasite was not the original focus and had never been found in bat fly samples before. One of the advantages of this technique is its capacity to uncover non-cultivable organisms, offering a more comprehensive view of parasite diversity in the environment.

To validate and deepen these findings, it is important to integrate other molecular tools, such as conventional PCR, and to attempt cultivation of less-studied species, followed by sequencing. This combined methodology helps not only to confirm the presence of the detected organisms, but also to generate important insights into their viability and biology, contributing to a better understanding of trypanosome ecology, especially in urban green areas.

*Trypanosoma wauwau*, first described in *Pteronotus parnellii* bats in Rondônia [10] and Pará [16], is the closest species to the sequences detected in this study. This trypanosome was found in three of the five fly samples parasitizing *Carollia perspicillata*, a frugivorous bat. Previous studies in the Atlantic Forest of Rio de Janeiro [17], Espírito Santo [18], and the Colombian Amazon [19] also reported trypanosomes associated with *C. perspicillata*. In SP, previously described species in bats include *T. cruzi* [12] and *T. madeirae* [20]. This is the first report of sequences from a Neobat4 clade in São Paulo state. Experimentally, *T. wauwau* showed no ability to infect mammals *in vitro*, mice, or triatomines, being inactivated in the latter’s digestive system [10].

In the phylogenetic analysis of the Neobat clades, the non-monophyletic distribution of the five distinct lineages (Neobat1–5) indicates an evolutionary history of the parasite marked by many independent events of adaptation to parasitism in bats. The bootstrap values provide variable support, since the Neobat2 clade (0.929) and its relationship with Neobat3 (0.899) stand out as well-supported taxonomic groups and Neobat3 displays a broad geographic distribution from Paraíba to Rio de Janeiro, suggesting a greater dispersal capacity or adaptability to different hosts. The Neobat4 clade exhibits a particular phylogenetic structure, with robust basal nodes (0.823 and 0.601) but low internal branching (0.003), indicating recent diversification events. Neobat1 and Neobat5 clades show distinct patterns, with Neobat1 presenting a close phylogenetic relationship with *Trypanosoma wauwau*, suggesting a possible shared evolutionary history and common ancestry.

Alves et al. [15] identified that *Carollia* sp. trypanosomes form a unique clade, Neobat4, possibly specific to this genus, a hypothesis reinforced by the sequences from this study.

The description of the Neobat clade of *Trypanosoma* sp. in Brazilian territory (São Paulo, Rio de Janeiro, Paraíba, Acre, and other unspecified locations) suggests a broader distribution of these parasites in South America, especially considering their presence in Acre state, which borders Peru and Bolivia. Further studies in other South American countries are needed to determine the true extent of this geographic distribution.

Regarding vectors, triatomines represent proven vectors, especially *Cavernicola* spp. [21], which preferentially feeds on bats in caves. In a study with *Cavernicola lenti*, protozoa with morphology similar to *T. cruzi marinkellei* was found [22]. In addition, in 1942, researchers documented *Cavernicola pilosa* and streblid flies collected from tree hollows that served as bat roosts, suggesting ongoing transmission [23]. More recently, *Triatoma vitticeps* carrying *Trypanosoma dionisii* was reported in Espírito Santo [18].

Beyond triatomines, experimental studies have shown that bedbugs of the species *Cimex lectularius* can act as possible vectors for some bat trypanosome species, such as *T. hedricki* and *T. myoti*. These bedbugs exhibit developmental patterns akin to those seen in *T. cruzi* within triatomines, which is especially significant given the high prevalence of these ectoparasites in urban bat roosts and their capacity to sustain transmission cycles [24].

The detection of *Trypanosoma* sequences exclusively in *Strebla guajiro* specimens associated with *Carollia perspicillata* reflects the complexity of parasite-host-vector interactions in green areas. This finding may be explained by cross-contamination through occasional insect ingestion by frugivorous bats [25], direct fly ectoparasitism, or poorly understood ecological interactions. The taxonomic distinction between *S. guajiro* (Streblinae) and *Trichobius joblingi* (Trichobiinae) may be relevant, considering their different parasitic behaviors: *S. guajiro* primarily inhabits and moves through the host’s fur, while *T. joblingi* is typically found on membrane surfaces.

Regarding clinical aspects, while trypanosome infections in bats are generally considered asymptomatic, the first case of clinical disease associated with trypanosomiasis in chiropterans was described and authors could observe hemolytic anemia, jaundice, and hemoglobinuric nephrosis [26]. Pathological examination revealed numerous organisms in blood and lymphoid tissues, along with mild interstitial pneumonia and hepatic sinusoidal leukocytosis, suggesting a systemic inflammatory process. Scientific evidence shows that bats have unique mechanisms to tolerate inflammation caused by viral infections [27], but the regulation of specific inflammatory responses to trypanosome infections remains poorly understood.

Data from 2011 obtained with *T. cruzi* strains isolated from urban bats showed infectivity potential for human cells, although with lower efficiency compared to established strains. Despite the low transmission risk, this finding indicates that surveillance of these parasites in bat populations remains a relevant public health concern [28].

Climate change and land use alterations can reshape ecological interactions between bats and their ectoparasites. Such shifts may contribute to the emergence of pathogen spillover and alter the dynamics of trypanosome transmission [29].

It is important to note that the scarcity of studies on bat trypanosomes, especially in Neobats clades, limits our ability to make more comprehensive interpretations.

Complementary characterization of the trypanosome diversity in bats is important to expand our understanding of these parasites’ evolution and ecology. Additional analyses using complementary molecular markers such as gGAPDH and Cytb and a broader geographic sampling are needed. Moreover, developing more sensitive molecular techniques for detecting uncultured species is essential, similar to metagenomic and metabarcoding approaches already successfully used in viral taxonomy, to better understand the evolutionary relationships within this protozoan group.

## CONCLUSION

This is the first report of *Trypanosoma* sp. detection in flies of the Streblidae family parasitizing *Carollia perspicillata* bats, a finding made possible by shotgun metagenomics. It is also the first report of a Neobat 4 clade *Trypanosoma* sp. in the Atlantic Forest of São Paulo state. The identification of this parasite–host– vector association reinforces the need for continued surveillance to better understand this parasitism’s dynamics, impact on bat health, as well as the potential risks to public health. Although there is no proof of Streblidae’s vector competence in *Trypanosoma* transmission, their close association with bat hosts suggests they may act as useful sentinels for detecting circulating pathogens, particularly in scenarios where direct sampling of bat hosts is not feasible, contributing to monitoring efforts with minimal disturbance to wildlife. Although additional studies are needed, particularly involving deeper molecular characterization and investigation of ecological interactions, this preliminary finding contributes to documenting the occurrence of this *Trypanosoma* sp. clade in streblid flies from the Atlantic Forest of São Paulo, expanding current knowledge on its geographic distribution.

## ACKNOWLEDGMENTS

To Dr. Izilda Curado, from Instituto Butantã, Dr. Carlos Lamas from the Zoology Museum of USP and to Dr. Juliana T. de Deus, from Sucen/Instituto Pasteur, for the support. To the other staff members and interns of São Paulo City Hall – Secretariat of Health and Environment who contributed to field work.

## AUTHOR CONTRIBUTIONS

**R.M**. conceived this study, field work, laboratory experiments, data analysis, and wrote the manuscript, **L.A.M.F**. manuscript revision and lab support, **E.C.R**. laboratory experiments, **M.S.N**., **E.T.B.C**., **A.R.R**., **D.C.O:** field work, logistics and data collection, **J.L.S**. manuscript revision, **G.G**. specimen identification and critical manuscript revision, **E.C.S**. supervised the study, revised the manuscript, and provided funding. All authors read and approved the final version of the manuscript.

The authors have no conflicts of interest to declare.

**Note:** AI-based language model was used to produce a preliminary English translation of the manuscript.

